# In vivo feasibility study of the use of porous polyhedral oligomeric silsesquioxane implants in partial laryngeal reconstruction

**DOI:** 10.1101/587691

**Authors:** Martin A. Birchall, Peggy Herrmann, Paul Sibbons

## Abstract

**Background:** Loss of substantial volumes of laryngeal tissue after trauma or cancer significantly impairs quality of life. We hypothesised that repair of laryngeal defects with a candidate biomaterial, seeded with mesenchymal stem cells (MSC) and epithelial cells, may offer a therapeutic approach to this unmet need.

**Method:** Moulded porous polyhedral oligomeric silsesquioxane polycarbonate-urea (POSS-PCU) scaffolds were seeded with human-derived MSC and epithelial cells, were implanted orthotopically into a defect created in the thyroid cartilage in eight pigs and monitored *in vivo* for 2 months. *In vivo* assessments were performed at 1, 2, 4 and 8 weeks post implantation. Histology was performed following termination.

**Results:** Implant operations were uncomplicated. One pig was terminated early (2 weeks post-implantation) following expectoration of its implant. No other mortality or morbidity was observed. Endoscopy showed partial extrusion of implants at two weeks and complete extrusion of all implants by termination.

**Conclusions:** POSS-PCU moulded laryngeal implants, in the present formulation, are extruded from the site of implantation between two- and eight-weeks post-surgery in pigs. In its present formulation and with the present, one-stage, protocol, this material does not appear to provide a suitable scaffold and vehicle for cells intended for partial laryngeal replacement in pigs.

## Introduction

Loss of substantial volumes of laryngeal tissue after trauma or cancer significantly impairs quality of life. Loss of a functioning larynx causes significant problems with speech, breathing, swallowing, and other core human activities.

Laryngeal allo-transplantations have been reported to date ^1,2,3^ but require life-long immunosuppression, with associated ethical and morbidity issues. Reinnervation, and thus control, of such transplants is also incomplete^4^. Tissue-engineering, using biologic or synthetic scaffold materials, is continuing to evolve and several tissue-engineered products are now available for clinical use^5^. In the airways, clinical success has recently been reported using both synthetic (polypropylene mesh with collagen sponge^6,7^) and biologic (homograft aorta^8,9^) scaffolds.

Most *in vivo* studies of laryngeal replacement are limited to the relatively small volume of the vocal cords^10,11^. A porcine derived multi-laminate extracellular matrix restored 70% of a larger defect with apparently good restoration of all tissue layers^12^.

Our previous reports showed feasibility^13^, safety and potential efficacy^14^ of a de-cellularised hemi-larynx seeded with human bone marrow derived mesenchymal stem cells (BM-MSC) in pigs. However, access to human donor larynxes may be a rate limiting supply chain feature of the clinical application of this approach. The provision of a biocompatible, customised biomaterial alternative, similarly seeded, might resolve this issue.

Porous polyhedral oligomeric silsesquioxane (POSS) are a candidate class of materials with cage-like structures at the molecular level with a surface that can be easily modified in various ways. POSS is widely used in industry, for example in circuit-printing^15^, but has also been modified for biomedical applications, such as copolymers for photodynamic therapy, controlled drug release and osteogenesis^16-19^. Of this family, POSS-PCU (POSS-polycarbonate-urea) appeared to have important candidate properties, specifically the ability to retain geometry *in vivo*^20^.

We hypothesised that repair of laryngeal defects with POSS-PCU implants, seeded with mesenchymal stem cells (MSC) and epithelial cells, would support structural regeneration of laryngeal tissue *in vivo* in pigs.

## Methods

### Ethics

Ethical permission was obtained from NPIMR Ethical Review Committee for animal experimentation and in accordance with the UK Animal (Scientific Procedures) Act 1986, which conforms to the European Convention for the protection of Vertebrate Animals Used for Experimental and Other Scientific Purposes (Strasbourg, Council of Europe).

### Scaffold Production

#### 1. Manufacture of 18% w/w polyhedral oligomeric silesquioxane poly (carbonate-urea) urethane (POSS-PCU) in dimethylacetamide (DMAC)

Polycarbonate polyol, and trans-cyclohexanechloroydrinisobutyl-silsesquioxane were placed in a reaction flask equipped with mechanical stirrer and nitrogen inlet. The mixture was heated to 135°C to dissolve the POSS cage into the polyol and then cooled to 70°C. Flake 4,4’-Methylenebis (phenyl isocyanate), (MDI), was added to the polyol blend and then reacted, under nitrogen, at 70°C - 80°C for 2 h to form a pre-polymer. Dimethylacetamide (DMAC) was added slowly to the pre-polymer to form a solution. The solution was cooled to 40°C. Chain extension of the pre-polymer was carried out by addition of Ethylenediamine in Dimethylacetamide to form a solution of POSS modified Polycarbonate urea-urethane in DMAC. All chemicals and reagents were purchased from Sigma-Aldrich (Gillingham, UK).

#### 2. Fabrication of non-porous POSS-PCU

POSS-PCU solution was poured on to a stainless-steel mould (diameter = 10 cm). To obtain 2 mm thickness of non-porous form. After pouring, the mould was placed in the incubator at 65° C and left overnight for at least 18 hours to allow the DMAC to fully evaporate. The solidified POSS-PCU sheet (2-mm thick) was detached from the mould and was cut into 3 cm × 6 cm. This piece was placed in the mould again and was poured on top with micro-porous POSS-PCU which will be described in the next step.

#### 3. Fabrication of microporous POSS-PCU on top of non-porous POSS-PCU

Microporous POSS-PCU was prepared using 18% POSS-PCU in DMAC solution mixed with 50% wt. pre-sieved NaHCO_3_ particles (100-150 µm in diameter). The compositions were mixed in the ARE-250 planetary centrifugal mixer (ThinkyUSA, Inc., California, USA) at 2,000 rpm mixing step and 1,500 rpm defoaming step. This mixture was then poured onto the mould which was pre-attached with non-porous POSS-PCU at the bottom (Figure 1). Finally, the mould was transferred into the deionised water to initiate the phase inversion/particulate-leaching to allow the solvent exchange and let DMAC and NaHCO_3_ micro-particles to slowly dissolve to form microporous part. The water was changed regularly up to 5 times to wash away excess solvent and NaHCO_3_.

**Figure 1.**
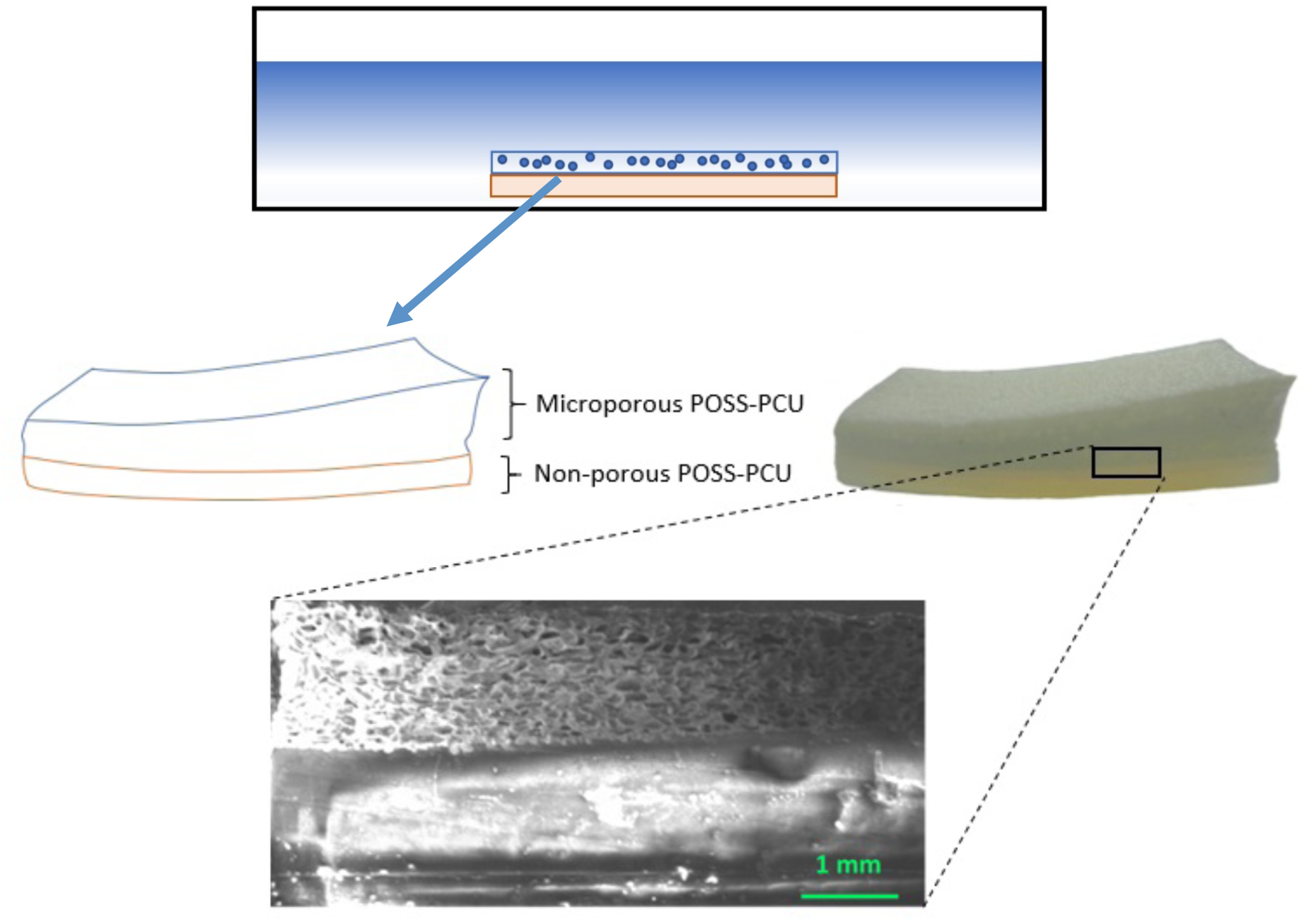
Schematic showing fabrication of laryngeal scaffold using combination of non-porous and microporous forms of POSS-PCU, with (lower panel) close up image of the interface between the two.

### Animals

The same species of animals was used as in previously published for our feasibility study using biologic scaffolds^13^. Briefly, eight female White/Landrace cross bred pigs, 45-50kg, were acclimatised for two weeks prior to the implantation surgery. Since xenogeneic cells were used, they were given 10-15mg/kg Cyclosporine (Neoral, Novartis, UK) commencing on the day before surgery, and then daily until termination. Interval endoscopy was performed before termination at two months.

### Cell culture and seeding

#### Human bone marrow derived mesenchymal stem cells (BM-MSC)

Again, the same approach was used as in our previously published feasibility study using biologic scaffolds^13^. In brief, human BM-MSC were obtained from consenting donors undergoing therapeutic MSC and cultured for three to seven passages in supplemented αMEM (Minimal Essential Medium, Lonza, USA). Hemi-larynges were seeded in dedicated bioreactors at a seeding density of 1.5×10^6^/cm^2^.

#### Primary Human airway epithelial cells

Human epithelial cells were prepared from small biopsies or resections of airway mucosa obtained with ethical approval and consent and expanded *ex-vivo* in minimum quantities of BEGM medium in a T25 flask and allowed to adhere overnight. After one week in an incubator (37°C, 5% CO_2_) flasks were topped up with medium and cultured for another week followed by washing, trypsinization, re-plating and expansion for three passages. Seeding onto the luminal aspect of each scaffold was 72h before surgery, with H&E staining of supplementary material used to confirm cell presence before release.

#### Scaffold seeding

Again, the same protocols were used for seeding as previously reported for seeding of biologic-based scaffolds^13^. In brief, prepared, sterilised scaffolds were place in Bronchial Epithelial Growth Medium (BEGM BulletKit, Lonza CC-3170, USA) in customised bioreactors. Seeding of epithelial cells on to the scaffold was carried out 72 hours before transplantation. Cells were prepared, trypsinized, counted and re-suspended in BEGM (epithelial cells) or αMEM (MSC). Epithelial cells were applied using a wide-tip pipette at 200,000 cells/cm^2^ and incubated for 48 hours, when the medium was changed, cells counted and MSCs seeded at a density of 1.5×10^6^cells/cm^2^. Seeded scaffolds were incubated for a further 24h before surgery.

### Surgery and in life assessment

Pigs (n=8) were pre-medicated with ketamine (Xylazine 1mg/kg Bayer UK, National Veterinary Service, Stoke-on-Trent, UK) intramuscularly. General anaesthesia (GA) was induced with isoflurane over oxygen and nitrous oxide. After intubation, anaesthesia was maintained with isoflurane. Cefuroxime 750mg (Flynn Pharma Ltd, Stevenage, UK) was given intravenously, subcutaneous analgesia Carprofen 4mg/kg (Rimadyl, Pfizer, Walton Oaks, UK) and subcutaneous anthelmintic (Ivermectin, Boehringer, Georgia, US) 1ml standard dose was given and Hypromellose eye drops (FDC Int Ltd, Fareham, UK).

Animals were randomised into groups for either left or right sided implantation using a computer-generated code. The larynx was exposed via a midline incision and retraction of the strap muscles. A musculo-perichondrial flap was raised on the appropriate side of the thyroid cartilage and a rectangular block of thyroid cartilage (2.5cm × 4.5cm) with associated tissue and ipsilateral vocal fold was removed. The implants were sutured in place with 4-0 Vicryl (Ethicon, Wokingham, UK) by direct suture to the thyrohyoid and cricothyroid membrane, as well as to the remaining thyroid cartilage. The perichondrial flap was repaired over the implant. Skin closure was in layers with 4-0 Vicryl and 2-0 Prolene, (both Ethicon, Wokingham, UK). Post-operative care is as previously described^13^.

### In life analysis

At weeks 1, 2, 4 and 8, endoscopy was performed under spontaneous ventilation, with cytology brushings of the implanted area and blood samples for haematology and clinical chemistry. CT scans were performed prior to terminal anaesthesia at week 8.

### Histology and Immuno-histochemical (IHC) analysis

Explanted larynges were photographed and fixed in 10% neutral buffered formal saline. Following fixation, each larynx was cut transversely at 0.5cm intervals and segments processed to paraffin wax blocks. 5 µm thickness samples were stained with Haematoxylin and Eosin stain (H&E).

Cytology brush samples were fixed, washed in PBS for 5min and non-specific binding sites were blocked using 10% FBS, PBS and 0.2% fish skin gelatine. Following incubation with primary monocolonal antibody (mouse anti-human cytokeratin 14, Abcam, Cambridge, UK) slides were washed and incubated with secondary antibody (red goat anti-mouse, Abcam, Cambridge, UK) in blocking solution. Finally, cell smears were stained with DAPI and washed. Human epithelial cells served as “positive” controls for human origin of retrieved cells, whilst porcine endolaryngeal cell smears acted as negative controls.

### Results

This was a feasibility study designed to determine whether a synthetic material POSS-PCU was 1) safe and 2) contributed to laryngeal regeneration following orthotopic implantation in a large animal model. Eight pigs received a cell-seeded POSS-PCU scaffold. All operations were uncomplicated. One pig was terminated early (2 weeks post-implantation) following expectoration of its implant. Otherwise, there was no morbidity or mortality and none required veterinary intervention due to dyspnoea, dysphagia or dysphonia. All animals had normal white blood cell and lymphocyte counts for the duration of the study.

#### Mucosal Cytology

Mucosal brushing taken at early time points (weeks 2 and 4) during bronchoscopy examination showed cells with epithelial cell morphology that stained positively for CK14, suggesting human origin. No phenotypically-human cells were detectable beyond four weeks.

#### Bronchoscopy

Bronchoscopy at 1, 2, 4 and 8 weeks’ post-implantation consistently showed a smooth appearance suggesting mucosal regeneration, ingress or draping. Representative images at each time point and for each scaffold type are presented in Figure 2.

**Figure 2.**
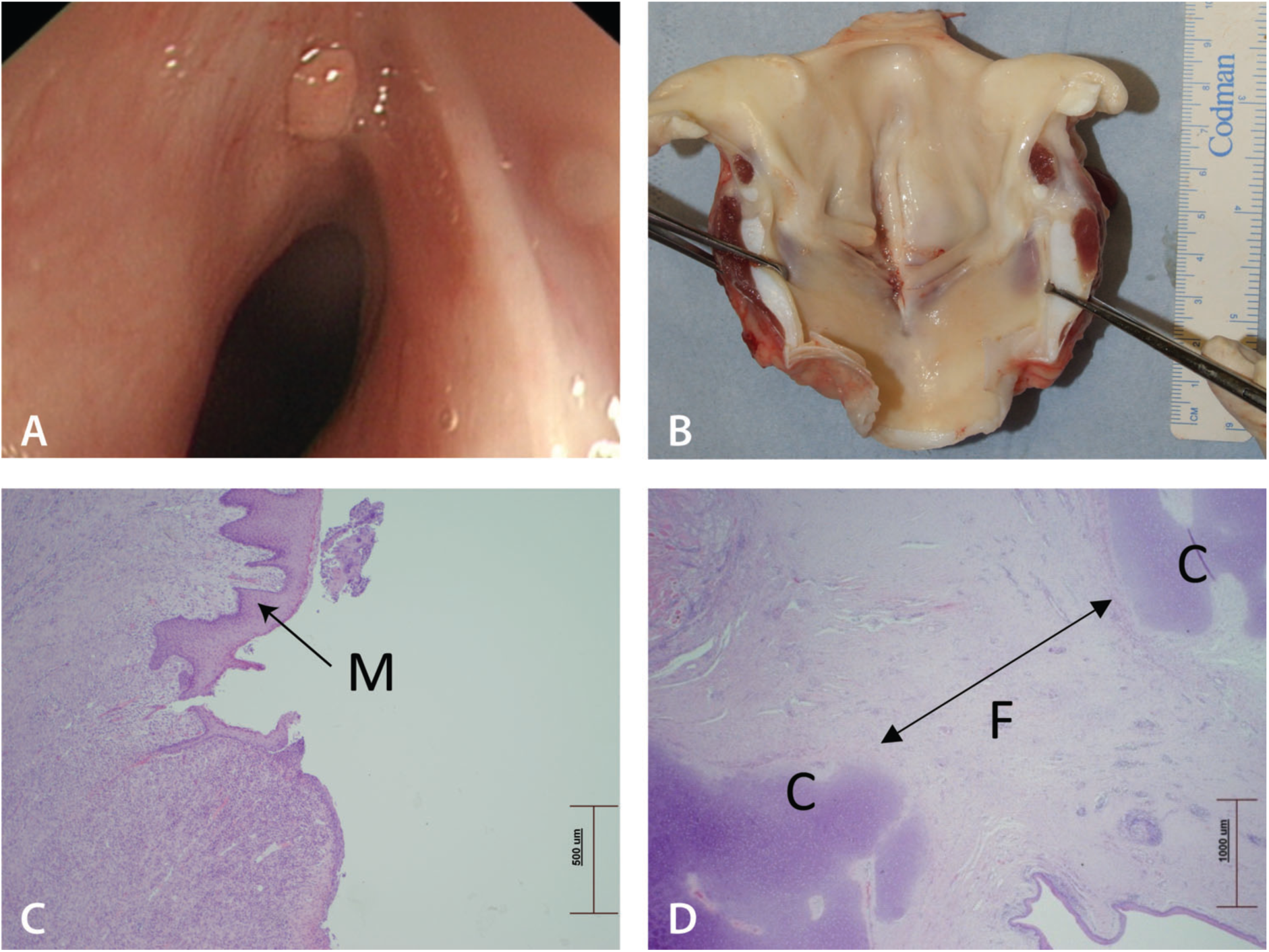
Representative bronchoscopy image showing a smooth mucosal covering (A) confirmed on explantation (B) and histological assessment (C&D), M= mucosal coverage C= host cartilage and F area of fibrosis.

#### Laryngeal explant analysis

On examining explanted larynges at termination, the internal mucosal surfaces all showed complete mucosal coverage: it was not possible to identify the surgical defect from the luminal surface. There was complete healing of the wound with little evidence of scar.

#### Histology

POSS-PCU material could not be detected in any sectioned larynx post-mortem. This suggested universal total extrusion. The luminal surface showed complete epithelialisation with rugi formation and mature vasculature. The cut edges of the cartilage could be identified and, between these, a mixed acute and chronic inflammatory response was observed. At high magnification, tracks of leukocytes surrounded by fibroblasts suggesting early fibrosis was observed.

### Discussion

We performed a feasibility study to test the hypothesis that repair of laryngeal defects with POSS-PCU implants, seeded with mesenchymal stem cells (MSC) and epithelial cells, would support structural regeneration of laryngeal tissue *in vivo* in pigs. Eight pigs received seeded implants. One expectorated its implant at two weeks and was terminated at that point, whilst all others survived to the censor point of eight weeks without any morbidity or mortality. Although, on termination, all animals alive at eight weeks showed complete mucosal coverage of the implanted area and normal phonation, breathing and swallowing, there was no sign of any of the implants in explanted larynges.

We have previously shown that tissue engineered partial laryngeal replacements based on recellularised biological (decellularised donor tissue) scaffolds are safe and encourage functional laryngeal regeneration in pigs^13,14^. However, preparation of clinical products based on biologic scaffolds requires a dependable supply chain of organs of varying size and gender from human donors, intensive sterilisation, and means of controlling reproducible and sustainable biomechanical strength. Thus, consideration of a synthetic scaffold source is reasonable.

POSS-PCU is a polymer that is widely used in biomedical applications^15-19^ and has characteristics that are potentially desirable in a laryngeal scaffold. Polycarbonate (urea urethane) is known to promote cell infiltration^21^ and, together with the POSS nanocage, was expected to form a suitable implant for partial laryngeal reconstruction.

It has been recently shown that these materials may be further modified to include a foam-like internal structure which could have implications towards long-term integration and reduced inflammation *in vivo* ^22^.

Although surgical implantation and post-operative course was uncomplicated in all animals in this study, and only one was terminated early due to expectoration of implant, at termination, none of the POSS-PCU implants had been retained. This suggests that, in the present formulation, this material does not integrate well into the laryngeal airway and is not, as it stands, a suitable scaffold for laryngeal tissue engineering.

We and others have used two-stage procedures with pre-implantation of constructs into well-vascularised areas such as the forearm or muscle, in order to pre-vascularise implants and support tissue ingress into implants for use in airway reconstruction^14,23^. It is possible that this strategy might be applicable to implants based on polymeric scaffolds also and might be a more successful approach than the present direct, one-stage, protocol if improved formulations of POSS-PCU, other POSS family polymers, or other biomaterials, are explored for this purpose in future.

In conclusion, POSS-PCU in the present formulation and with the present protocol does not appear to be a suitable material for partial laryngeal replacement in pigs.

## Acknowledgements

Assistance with fabrication of POSS PCU was given by Mr Arnold Darbyshire, Division of Surgery, UCL. The authors particularly would like to express their gratitude for the professional assistance given by the theatre and husbandry team at NPIMR during the surgical procedure and after-care of pigs during this study.

